# *In vivo* human uterine temperature, pH, and uterine fluid composition analysis

**DOI:** 10.1101/2024.11.18.623470

**Authors:** M. Tarahomi, M.S. Zagers, S. Zafardoust, A. Mohammadzadeh, Z. Fathi, H. Sareban, F. Fatemi, S. Fakhr, G. Hamer, S. Repping, F.A.P. Schrauwen, J. P. van Straalen, F.M. Vaz, M. van Wely, S. Mastenbroek

## Abstract

**Study question:** What are the temperature, pH and uterine fluid composition in the human uterus three days following a positive LH test or ovum pick-up?

**Summary answer:** The mean uterine temperature was 36.94±0.26°C, the mean uterine pH was 6.76±0.22, and the concentrations of 37 components in aspirated uterine fluid were successfully determined.

**What is known already:** Embryo culture conditions in the laboratory impact key outcomes of IVF/ICSI treatments, such as the quality of the embryos and the live birth rate after treatment, and child outcomes, such as birth weight. Currently used conditions, including temperature, pH, and culture medium composition, are largely derived from clinical experience and experimental studies using animal models. Limited studies have been performed to determine the natural human preimplantation embryo environment *in vivo* during the physiologically relevant time of the menstrual cycle. This type of fundamental knowledge is required for evidence-based optimization of the *in vitro* embryo culture environment and improving IVF/ICSI outcomes.

**Study design, size, duration:** In this cross-sectional study, conducted between April 2015 and March 2016, temperature and pH were measured in the human uterine cavity on the third day following a positive LH test or ovum pick-up, and uterine fluid was simultaneously aspirated for composition analysis. Uterine temperature was measured in fifty eight women, uterine pH was determined in fifty three women, and twenty two samples of aspirated uterine fluid were analysed for the concentrations of thirty-seven components.

**Participants/materials, setting, methods:** This study involved 61 healthy reproductive-aged women: 53 without ovarian stimulation and 8 who underwent ovarian stimulation. We measured uterine temperature using a probe inserted into the uterine cavity directly, and uterine pH after inserting a probe through the outer sheath of an IVF catheter. Uterine fluid was then aspirated using this outer IVF catheter and a 10 ml syringe, and subsequently analysed with a Cobas 8000 chemistry analyser and ultra-performance liquid chromatography-tandem mass spectrometry.

**Main results and the role of chance:** The mean uterine temperature on the third day following a positive LH test or ovum pick-up was 36.94 ± 0.26°C and correlated with the women’s core body temperature. The mean pH in the uterine cavity was pH 6.76 ± 0.22, clearly lower than the standard pH used for human preimplantation embryo culture *in vitro* (pH 7.3 ± 0.1). Concentrations of important energy sources were 0.8 ± 0.02 mM pyruvate, 5.1 ± 1.78 mM glucose and 6.60 ± 1.12 mM lactate. Glutamic acid (1162 ± 183 μM), glycine (955 ± 156 μM) and alanine (513 ± 82 μM) were the most abundant amino acids in uterine fluid.

**Limitations, reasons for caution:** In absence of a preimplantation embryo, synergistic influences on the uterine environment may be overlooked. Single centre and specific population limitations may hinder broader generalization of the results. Uterine fluid likely contains additional components.

**Wider implications of the findings:** The *in vivo* uterine characteristics identified in this study are foundational to develop an *in vivo* evidence-based culture medium for human embryos. Further research is necessary to evaluate whether such a medium can improve human preimplantation embryo development and quality, thereby increasing cumulative live birth rates and improving child outcomes.

## Introduction

Since the first live birth following IVF in 1978, more than ten million babies have been born worldwide through IVF (ESHRE, 2023). Despite the large number of live births, the IVF success rates remain suboptimal, with approximately two-thirds of treatment cycles not leading to a successful pregnancy. In 2019, the cumulative live birth rate was reported to be 30.7% (ESHRE, 2023). A potential limiting factor may be a suboptimal *in vitro* culture environment that is not fully equipped to support human preimplantation embryo development (Chronopoulou and Harper, 2015). Primary elements of this environment, including CO_2_/O_2_ levels, temperature and the composition, pH and osmolality of the culture medium, influence the quality of preimplantation embryos – a key predictor for the chance of a live birth (Feuer and Rinaudo, 2012, Swain, 2015, Gardner and Kelley, 2017, Cairo Consensus Group, 2020). Comparative clinical trials on human embryo culture media have demonstrated that the choice of embryo culture medium affects cumulative live birth rates and child outcomes, such as birth weight (Dumoulin et al., 2010, Nelissen et al., 2012, Youssef et al., 2015, Kleijkers et al., 2016). Optimizing the *in vitro* embryo culture environment may thus improve preimplantation embryo development and quality, and subsequently increase the chances of a live birth and improve child outcomes.

Current human embryo culture conditions primarily rely on laboratory experience and experimental studies with animal models on the effects of altering culture parameters on embryo development and/or IVF outcomes. Most of these studies have been conducted with mouse embryos or other animal models, due to the fact that human preimplantation embryos are generally unavailable. The mouse embryo assay is often used for development and testing of commercial human embryo culture media. However, designing and optimizing a human preimplantation embryo culture environment based on experimental mouse or other animal embryo studies may lead to a culture system tailored to mouse preimplantation embryos that is less applicable to human preimplantation embryos (Menezo and Herubel, 2002, Chronopoulou and Harper, 2015). Given that embryo culture media consist of various components with interactive effects, obtaining empirical evidence would also require examining potentially endless possibilities (Summers and Biggers, 2003). Even if human embryos had been available, achieving a truly optimized human preimplantation embryo culture environment following an approach of studying individual culture medium components or culture parameters alone, therefore seems implausible (Summers and Biggers, 2003, Pool et al., 2012, Chronopoulou and Harper, 2015). A “back to nature approach” – striving to mimic the natural *in vivo* human preimplantation embryo environment *in vitro* (Leese, 1998) – has emerged as essential for achieving true optimization. This approach requires a comprehensive understanding of the *in vivo* human preimplantation embryo environment. However, limited studies have been performed to elucidate the *in vivo* characteristics of the preimplantation embryo environment in humans, such as temperature, pH and uterine fluid composition.

### Temperature

The commonly used temperature for human embryo culture in IVF laboratories is 37.0°C and is based on the estimated core body temperature (Swain, 2015). Observations in animal models and a study measuring human follicle fluid temperature 2.3°C lower than the ovarian stroma, suggested optimization of *in vitro* culture by (partially) culturing human embryos at temperatures below 37.0°C (Grinsted et al., 1985). But studies investigating the impact of human embryo culture at lower or gradual temperatures failed to yield conclusive results (Baak et al., 2019). Further *in vivo* evidence on the temperature in the human reproductive tract is limited to a single study reporting a uterine temperature of 37.08 ± 0.18°C in the ovulatory phase (n = 10) and 37.47 ± 0.35°C in the secretory phase (n = 11) (Yedwab et al., 1976). A limitation of this study is its reliance on alterations in core body temperature to estimate menstrual cycle phases, introducing uncertainty regarding the specific timing of temperature measurements during the menstrual cycle. Also, a limited number of measurements were performed and temperature meter accuracy has likely been improved since then. Further research into the temperature in the human reproductive tract seems required to determine the temperature preimplantation embryos naturally develop in.

### pH

Embryologists generally culture human embryos within a range of pH 7.2-7.4. Commercial embryo culture media manufacturers commonly recommend this pH range in the instructions accompanying the embryo culture media. The current standard pH of 7.2-7.4 appears to originate from the initial use of a culture medium with a stable pH of 7.6 developed for hamster embryos, for the culture of human gametes to enable fertilization, and further refinement lowering the pH through studies on different culture media, pH, and bicarbonate buffering conditions for human embryo culture (Edwards and Steptoe, 1974, Edwards et al., 1981, Bavister, 2002). The embryo culture medium pH is important as it influences the intracellular pH (pHi), impacting cellular processes with potential consequences for embryo development (Busa and Nuccitelli, 1984, Squirrell et al., 2001, Swain and Pool, 2009, Gu et al., 2023). Induced changes in the culture medium pH for a short period during embryo culture have been shown to affect the birthweight of mouse pups (Banrezes et al., 2011). Despite its importance, the optimal pH for the human preimplantation embryo environment remains to be determined (Swain, 2010, Swain, 2012, Gatimel et al., 2020). Only three studies have previously measured the pH in the human reproductive tract, all reporting pH values lower than the pH range of 7.2-7.4 that is commonly used for *in vitro* human embryo culture worldwide (Feo, 1955, Sedlis et al., 1967, Yedwab, et al., 1976). pH measurements in these studies were conducted on different days throughout the menstrual cycle and did therefore only partly represent the natural environment of a preimplantation embryo. Additional *in vivo* pH measurements are required to identify the pH of the natural preimplantation embryo environment and to assess the suitability of the pH range currently used for *in vitro* human embryo culture.

### Oviduct and uterine fluid composition

Many different human preimplantation embryo culture media are commercially available and globally used in IVF laboratories. The varying concentrations of components across brands and types of media indicate that the optimal composition for human preimplantation embryo culture medium still remains unknown (Morbeck et al., 2014, Morbeck et al., 2017, Tarahomi et al., 2019, Zagers et al., 2023). While some of these concentrations seem to be based on limited available evidence on the composition of human oviduct and uterine fluid, most concentrations appear to be derived from experimental findings (Morbeck, et al., 2014, Morbeck, et al., 2017, Tarahomi, et al., 2019, Zagers, et al., 2023). Sixteen studies have previously determined the concentrations of several or more components of human oviduct or uterine fluid (Lippes et al., 1972, David et al., 1973, Lippes, 1975, Shams et al., 1977, Borland et al., 1980, Casslen and Nilsson, 1984, Dickens et al., 1995, Gardner et al., 1996, Srivastava et al., 1996, Tay et al., 1997, Chen et al., 2002, Strandell et al., 2004, Boomsma et al., 2009, Hannan et al., 2011, Kermack et al., 2015, Utsunomiya et al., 2022). Most of these studies aspirated oviduct or uterine fluid throughout the menstrual cycle instead of specifically within a few days after ovulation – the time a preimplantation embryo would reside in the oviduct or uterus. Other limitations were obtaining fluid from the oviduct or uterus *ex vivo*, analysing hydrosalpinx fluid, collecting oviduct fluid after vascular perfusion, analysing samples of low volume, analysing a limited amount of samples, or aspirating uterine or oviduct fluid from women who were either suspected of or diagnosed with uterine diseases and infertility. Eleven other studies focused on identifying proteins present in human oviduct fluid and/or human uterine fluid (Moghissi, 1970, Lippes et al., 1981, Lippes et al., 1983, Lippes and Wagh, 1989, Wagh and Lippes, 1989, Lippes and Wagh, 1993, Wagh and Lippes, 1993, Parmar et al., 2008, Salamonsen et al., 2013, Canha-Gouveia et al., 2019, Fujii et al., 2021). Nevertheless, a thorough understanding of the composition of human oviduct and uterine fluid is still lacking. Reliable oviduct and uterine fluid analyses from healthy women of reproductive age are required to ascertain the composition during the menstrual cycle phase when preimplantation embryos naturally reside in the oviducts and uterus.

To summarize, the *in vivo* evidence underlying the current culture conditions for human preimplantation embryos is scarce. A better understanding of the *in vivo* preimplantation embryo environment in humans is essential to enable mimicking of the natural embryo environment for optimization of *in vitro* human embryo culture conditions. Therefore, we measured the temperature and pH in the human uterine cavity on the third day following a positive LH test or ovum pick-up, and simultaneously aspirated human uterine fluid to analyse the concentrations of thirty-seven components.

## Materials and Methods

### Ethical approval

This study was conducted according to the principles of the Declaration of Helsinki (Oct 2013). Ethical approval was received in 2015 from the Ethics Committee of the Avicenna Research Institute (ARI), Teheran, Iran (decision number 258-15-95).

### Study population

Women between 18 to 43 years old with a regular ovulatory cycle and a normal uterine function who were visiting the Avicenna Infertility Clinic, Teheran, Iran, for fertility treatment due to severe male factor subfertility (azoospermia or a total motile sperm count lower than eight hundred thousand per ml of semen) or for PGT-M or sex selection treatment, were asked to participate in this study. Women with a history of HIV or any sexually transmitted disease, malformation of the urogenital system, borderline or invasive ovarian cancer or any other cancer, a history of dilatation and curettage, premature ovarian failure, PCOS or severe psychopathology were not eligible for this study. All women were informed about the study during their visit to the clinic and received an informative letter about the study and their personal fertility treatment. In total, sixty-one women provided written informed consent to participate in this study between April 2015 and March 2016. Fifty three women participated during a natural menstrual cycle before the start of their planned fertility treatment. Eight women participated during an ovarian stimulated cycle during their fertility treatment, three days after the ovum pick-up. Participating in this study during these stimulated cycles was possible because the freeze all embryos strategy was applied in these fertility treatments. Seven women did not receive a fresh embryo transfer due to ovarian hyperstimulation syndrome (OHSS) and one woman opted for elective freeze all.

### Timing in the menstrual cycle

To determine the time of ovulation in an unstimulated menstrual cycle, participating women were trained for daily utilization of the “Ovulation Prediction Kit” (OPK) (Sensitest). The OPK is a urine dipstick kit to ascertain the time of the LH surge, which predicts ovulation. Participants started testing on day nine of the menstrual cycle and tested daily until a test was positive. The women were asked to pay an extra visit to the clinic three days after the positive reading of the ovulation test for the uterine measurements and uterine fluid collection. The eight women participating during a stimulated cycle payed an extra visit to the clinic for the uterine measurements and uterine fluid collection on the third day following the ovum pick up.

### Uterine temperature and pH measurements

All women were positioned in lithotomic position and examined in the ovum pick-up room. Ultrasound imaging was used to evaluate the follicles, the pattern of the ovaries, possible cysts, the length of the uterus (distance between external orifice and fundus), the anterior to posterior uterine diameter, possible uterine malformations, endometrial aspects and thickness.

Without any cleansing of the vagina, a medically approved disposable general purpose temperature probe (81-020409EU; DeRoyal), connected to a general vital signs monitor, was then inserted in the posterior vaginal fornix to record the body temperature. We selected this probe because of the following characteristics described by the supplier: 1) its versatile design allows for oral, nasal or rectal introduction for routine monitoring of patient core body temperature, 2) it has a low-friction surface for ease of insertion, 3) it has a completely enclosed sensor wires for improved stability and hygiene,

4) the diameter is very thin (ø = 3mm) and its flexible design ensured optimal patient safety, 5) it is fully compatible with most monitors, 6) it was not made with natural rubber latex, 7) it is sterile and 8) it has a short response time of only 60 seconds. After this, the temperature probe was inserted through the cervix into the uterus following the standard procedure of a routine embryo transfer. To prevent any contact with the fundus or bleeding of the endometrium, the temperature probe was inserted into the uterine cavity up to four to five centimeters from the external orifice of the cervix. After removing the temperature probe from the uterine cavity, it was checked for the presence of blood on the tip.

The standard procedure for embryo transfer (ET) was then applied for insertion of the outer sheath of an ET catheter (IVFETFLEX). This was followed by insertion of a calibrated micro pH probe (pHersaflex S1I catheter; Medical Measurement System Company (MMS)/ Laborie) through this outer sheath of the ET catheter into the uterine cavity to one cm beyond the end of the outer sheath of the ET catheter and up to four to five centimeters from the external orifice of the cervix. This micro pH probe is a flexible, medically approved disposable micro probe with a diameter of 1.67 mm and a response time of 30 seconds at the latest. Therefore, the pH was measured and recorded every second until 60 seconds after insertion in the uterine cavity. The pH probe was connected to an Orion II pH recorder and used with MMS Software for calibration of each pH probe and the pH measurement. Calibration of each pH probe was performed with two known pH solutions in sachets (pH = 4 and 7; Mettler Toledo BV) prior to each measurement, following the company’s instructions. After removing the pH probe from the uterine cavity, it was checked for the presence of blood on the tip.

### Uterine fluid aspiration

Directly after pH measurement, a five ml syringe was connected to the outer end of the outer sheath of the ET catheter. Aspiration of uterine fluid was performed gently by applying a negative pressure with the syringe and withdrawing the outer sheath of the ET catheter from the uterine cavity simultaneously. Aspirated, often highly viscous, uterine fluid was immediately expelled from the outer sheath of the ET catheter into a 500 μl Eppendorf micro-tube using positive air pressure supplied through the syringe. The sample was spun down at 4000 rpm for 30 minutes at 37°C in an Eppendorf 5810R centrifuge to separate possible aspirated endometrial cells from the uterine fluid. Supernatant and pellet were snap frozen separately and stored in -80°C until composition analysis.

### Uterine fluid analysis

On the day of composition analysis, twenty-two collected uterine fluid samples (supernatants) were thawed and the volume of every sample was determined. To decrease the viscosity, the volume of each sample was increased up to 300 μl by adding Milli-Q water (Millipore) and mechanically shaken (1800 oscillations per minute for two minutes using a Tissue lyser II (Qiagen)). Concentrations of ions (calcium, chloride, iron, magnesium, sodium, phosphate, potassium), glucose, uric acid, immunoglobulins (IgA, IgG and IgM), albumin and total protein were measured using a Cobas 8000 chemistry analyser (Roche Diagnostics). Concentrations of lactate (L and D), pyruvate and 21 amino acids (alanine, arginine, asparagine, aspartic acid, citrulline, glutamic acid, glutamine, glycine, histamine, isoleucine, leucine, lysine, methionine, ornithine, phenylalanine, proline, serine, threonine, tryptophan, tyrosine and valine) were determined using Ultra-Performance Liquid Chromatography tandem Mass Spectrometry (UPLC-MS/MS; Acquity-Quattro Premier XE, Waters, Milford, Massachusetts, USA). Iron was measured in micromolar (μM) and all other components were measured in millimolar (mM). After composition analysis, the measured concentrations were recalculated to the original volume of the uterine fluid sample. All measurements were performed at once by two technicians who were blinded to the source and characteristics of each sample. To achieve this, coded labelling was applied for all samples.

### Statistical analysis

The pH and temperature determined in the uterine cavity are presented as the mean ± Standard Deviation (SD). The concentrations of components of uterine fluid are presented as the mean ± Standard Error of the Mean (SEM). To test whether there was a correlation between core body temperature and uterine temperature we performed a Pearson’s correlation test in IBM SPSS Statistics 28. To determine whether uterine temperature and pH significantly differed between unstimulated and stimulated women, we performed independent t-tests in IBM SPSS Statistics 28.

## Results

One hundred and fifty nine women attending the Avicenna Infertility Clinic (Tehran, Iran) for IVF/ ICSI treatment due to severe male factor subfertility or for PGT-M or sex selection were asked to participate in this study between April 2015 and March 2016. In total, one hundred and eight women signed the informed consent form. Eleven participants withdrew from participation before the appointment for uterine measurements and fluid aspiration. Thirty six participants were excluded due to absence of a positive reading of the LH tests or because they did not show up for the appointment due to travel difficulties from other cities or regions. Altogether, sixty one participants came to the clinic for uterine temperature and pH measurements and uterine fluid aspiration on the third day following a positive LH test (n = 53) or ovum pick-up (n = 8) (Table 1). Fifty three participants did not receive ovarian stimulation and were examined during their natural menstrual cycle before the start of their personal IVF/ICSI treatment. Eight participants did receive ovarian stimulation for IVF/ICSI treatment, but did not receive an embryo transfer in the same cycle as the ovum pick up due to ovarian hyper-stimulation syndrome (OHSS; participants 14, 18, 20–22, 26 and 34) or elective freeze-all (participant 16), and their uterine environment was examined instead. All participants were between 20 and 40 years old, with a mean age of 29.9 ± 4.4 years old. The mean BMI was 24.8 ± 3.3 kg/m2, within a range of 17.2-32.8 kg/m2. Nine women have been proven to be fertile, as they delivered one or two times before.

**Table 1.** Participants’ characteristics, including measured uterine temperature and pH. In the last row we presented the mean ± SD for age, BMI, vaginal temperature, uterine temperature and uterine pH. The vaginal temperature of participants 12 and 23 was not included in the calculation of the mean ± SD.

| Participant study number | Age | BMI | Previous gravidity | Semen analysis result partner | Hormonal stimulation for IVF treatment | Timing of uterine measurements and uterine fluid aspiration | Endometrial thickness (mm) | Endometrial pattern (layers) | Core body temperature (°C) | Uterine temperature (°C) | Uterine pH |
| --- | --- | --- | --- | --- | --- | --- | --- | --- | --- | --- | --- |
| 1 | 33 | 23,8 | 0 | Azoospermia | No | 3rd day following + LH test | 8,2 | C | 37,3 | 37,1 | 6,71 ± 0,02 |
| 2 | 24 | 29,4 | 0 | Oligospermia | No | 3rd day following + LH test | 6,7 | C | 37,3 | 36,9 | 6,39 ± 0,09 |
| 3 | 29 | 25,4 | 0 | Azoospermia | No | 3rd day following + LH test | 7,1 | C | 37,1 | 37,1 | 6,58 ± 0,05 |
| 4 | 31 | 24,0 | 2 | Normospermia | No | 3rd day following + LH test | 7,1 | C | 37,5 | 37,2 | 6,81 ± 0,05 |
| 5 | 33 | 24,8 | 0 | Azoospermia | No | 3rd day following + LH test | 11,8 | B | 37,1 | 36,9 | 6,75 ± 0,11 |
| 6 | 33 | 24,3 | 0 | Azoospermia | No | 3rd day following + LH test | 8,2 | C | 37,1 | 36,9 | 6,72 ± 0,04 |
| 7 | 27 | 24,7 | 0 | Azoospermia | No | 3rd day following + LH test | 8,2 | C | 37,1 | 36,6 | 6,80 ± 0,09 |
| 8 | 31 | 20,2 | 0 | Azoospermia | No | 3rd day following + LH test | 6,3 | C | 37,1 | 37,3 | 6,72 ± 0,18 |
| 9 | 33 | 23,4 | 0 | Azoospermia | No | 3rd day following + LH test | 12,6 | B | 37,0 | 36,8 | 6,59 ± 0,05 |
| 10 | 29 | 27,6 | 0 | Azoospermia | No | 3rd day following + LH test | 8,8 | B | 37,3 | 37,0 | n.a. |
| 11 | 31 | 30,4 | 0 | Azoospermia | No | 3rd day following + LH test | 8,3 | B | 37,0 | 37,1 | 6,83 ± 0,07 |
| 12 | 30 | 24,0 | 0 | Azoospermia | No | 3rd day following + LH test | 10,3 | B | 37,9 | n.a. | n.a. |
| 13 | 29 | 25,2 | 0 | Azoospermia | No | 3rd day following + LH test | 7,7 | B | 36,9 | 36,6 | n.a. |
| 14 | 30 | 22,5 | 0 | Azoospermia | Yes (OHSS) | 3rd day following ovum pick-up | 9,4 | A | 37,0 | 37,0 | 7,03 ± 0,07 |
| 15 | 25 | 26,4 | 0 | Oligospermia | No | 3rd day following + LH test | 8,4 | B | 37,1 | 36,8 | 6,72 ± 0,16 |
| 16 | 35 | 25,2 | 1 | Normospermia | Yes (elective freeze-all) | 3rd day following ovum pick-up | 8,4 | B | 37,1 | 37,0 | 6,95 ± 0,06 |
| 17 | 31 | 27,1 | 0 | Oligospermia | No | 3rd day following + LH test | 6,8 | B | 37,4 | 37,1 | 6,65 ± 0,07 |
| 18 | 25 | 20,7 | 0 | Azoospermia | Yes (OHSS) | 3rd day following ovum pick-up | 9,1 | A | 36,9 | 37,0 | 6,73 ± 0,08 |
| 19 | 37 | 27,5 | 2 | Normospermia | No | 3rd day following + LH test | 8,8 | C | 36,8 | 36,9 | 6,70 ± 0,03 |
| 20 | 32 | 28,5 | 0 | Azoospermia | Yes (OHSS) | 3rd day following ovum pick-up | 8,1 | A | 37,0 | 37,0 | 6,74 ± 0,17 |
| 21 | 35 | 32,8 | 2 | Normospermia | Yes (OHSS) | 3rd day following ovum pick-up | 10,5 | B | 37,0 | 37,0 | 6,79 ± 0,03 |
| 22 | 33 | 21,3 | 0 | Azoospermia | Yes (OHSS) | 3rd day following ovum pick-up | 8,8 | B | 37,3 | 37,2 | 7,00 ± 0,07 |
| 23 | 23 | 26,1 | 0 | Azoospermia | No | 3rd day following + LH test | 7,6 | C | 37,6 | n.a. | n.a. |
| 24 | 38 | 29,7 | 0 | Azoospermia | No | 3rd day following + LH test | 7,4 | C | 36,9 | 36,6 | 6,86 ± 0,06 |
| 25 | 27 | 26,4 | 0 | Azoospermia | No | 3rd day following + LH test | 10,1 | C | 37,3 | 37,0 | n.a. |
| 26 | 26 | 31,9 | 0 | Azoospermia | Yes (OHSS) | 3rd day following ovum pick-up | 6,2 | A | 37,2 | 36,7 | 6,53 ± 0,05 |
| 27 | 24 | 21,2 | 0 | Azoospermia | No | 3rd day following + LH test | 12,1 | B | 37,2 | 36,8 | 6,70 ± 0,00 |
| 28 | 23 | 20,2 | 0 | Azoospermia | No | 3rd day following + LH test | 8,2 | C | 37,2 | 36,8 | 6,95 ± 0,07 |
| 29 | 34 | 24,6 | 0 | Azoospermia | No | 3rd day following + LH test | 7 | C | 36,7 | 36,8 | 6,75 ± 0,13 |
| 30 | 27 | 25,0 | 0 | Oligospermia | No | 3rd day following + LH test | 6 | C | 36,8 | 36,8 | 6,60 ± 0,14 |
| 31 | 28 | 21,0 | 0 | Oligospermia | No | 3rd day following + LH test | 8 | C | 36,7 | 36,8 | n.a. |
| 32 | 31 | 26,0 | 0 | Oligospermia | No | 3rd day following + LH test | 10 | B | 37,3 | 37,3 | 6,97 ± 0,10 |
| 33 | 31 | 20,6 | 0 | Oligospermia | No | 3rd day following + LH test | 9 | B | 37,2 | 37,3 | 6,66 ± 0,27 |
| 34 | 35 | 27,8 | 0 | Azoospermia | Yes (OHSS) | 3rd day following ovum pick-up | 5,3 | C | 37,1 | n.a. | n.a. |
| 35 | 28 | 29,8 | 0 | Azoospermia | No | 3rd day following + LH test | 10,5 | B | 37,2 | 37,1 | 6,86 ± 0,15 |
| 36 | 24 | 22,6 | 0 | Azoospermia | No | 3rd day following + LH test | 6,7 | B | 36,3 | 36,6 | 6,59 ± 0,03 |
| 37 | 27 | 17,2 | 0 | Azoospermia | No | 3rd day following + LH test | 9,6 | B | 36,7 | 37,3 | 6,60 ± 0,03 |
| 38 | 29 | 27,1 | 0 | Azoospermia | No | 3rd day following + LH test | 8,8 | A | 36,7 | 37,1 | n.a. |
| 39 | 32 | 21,1 | 0 | Azoospermia | No | 3rd day following + LH test | 6,1 | C | 37,0 | 36,4 | 7,08 ± 0,04 |
| 40 | 29 | 28,6 | 0 | Azoospermia | No | 3rd day following + LH test | 7,3 | C | 37,0 | 36,8 | 6,76 ± 0,12 |
| 41 | 34 | 23,5 | 2 | Normospermia | No | 3rd day following + LH test | 11,7 | C | 37,1 | 37,1 | 6,50 ± 0,03 |
| 42 | 35 | 23,4 | 0 | Oligospermia | No | 3rd day following + LH test | 5,8 | C | 36,8 | 37,0 | 6,44 ± 0,05 |
| 43 | 29 | 21,4 | 0 | Azoospermia | No | 3rd day following + LH test | 13,2 | B | 37,0 | 36,9 | 6,99 ± 0,03 |
| 44 | 40 | 24,0 | 0 | Azoospermia | No | 3rd day following + LH test | 11 | C | 37,0 | 36,9 | 6,78 ± 0,04 |
| 45 | 27 | 28,0 | 0 | Oligospermia | No | 3rd day following + LH test | 12 | B | 36,9 | 36,8 | 6,72 ± 0,04 |
| 46 | 21 | 22,8 | 0 | Azoospermia | No | 3rd day following + LH test | 10,2 | A | 37,4 | 37,0 | 6,60 ± 0,00 |
| 47 | 24 | 22,0 | 0 | Azoospermia | No | 3rd day following + LH test | 7 | C | 37,0 | 36,9 | 7,25 ± 0,09 |
| 48 | 21 | 25,3 | 0 | Oligospermia | No | 3rd day following + LH test | 6,6 | C | 37,0 | 37,0 | 6,48 ± 0,06 |
| 49 | 34 | 17,9 | 0 | Azoospermia | No | 3rd day following + LH test | 11,9 | C | 36,9 | 36,8 | 6,93 ± 0,08 |
| 50 | 36 | 21,1 | 0 | Azoospermia | No | 3rd day following + LH test | 5,9 | C | 37,5 | 37,1 | 6,73 ± 0,11 |
| 51 | 36 | 27,2 | 0 | Azoospermia | No | 3rd day following + LH test | 7,9 | C | 36,9 | 37,1 | 7,21 ± 0,11 |
| 52 | 34 | 25,7 | 0 | Azoospermia | No | 3rd day following + LH test | 5,6 | C | 37,1 | 37,0 | 7,05 ± 0,05 |
| 53 | 31 | 27,7 | 1 | Azoospermia | No | 3rd day following + LH test | - | - | 36,7 | 36,9 | 6,83 ± 0,12 |
| 54 | 28 | 24,0 | 0 | Azoospermia | No | 3rd day following + LH test | 7,6 | A | 37,0 | 37,0 | 6,87 ± 0,11 |
| 55 | 28 | 19,5 | 0 | Azoospermia | No | 3rd day following + LH test | 9,4 | C | 37,0 | 36,7 | 6,31 ± 0,04 |
| 56 | 20 | 24,5 | 0 | Azoospermia | No | 3rd day following + LH test | 10 | C | 36,9 | 36,0 | 6,91 ± 0,19 |
| 57 | 30 | 28,7 | 1 | Normospermia | No | 3rd day following + LH test | 10 | C | 37,2 | 36,9 | 6,59 ± 0,03 |
| 58 | 34 | 28,4 | 1 | Normospermia | No | 3rd day following + LH test | 9 | C | 36,9 | 36,6 | 6,50 ± 0,02 |
| 59 | 29 | 22,0 | 0 | Azoospermia | No | 3rd day following + LH test | 6,7 | B | 37,0 | 37,0 | 7,34 ± 0,06 |
| 60 | 30 | 25,6 | 1 | Normospermia | No | 3rd day following + LH test | 12,2 | B | 37,4 | 37,6 | 6,39 ± 0,11 |
| 61 | 32 | 24,4 | 0 | Azoospermia | No | 3rd day following + LH test | 8,2 | B | 37,5 | 37,6 | 6,81 ± 0,06 |
| Mean ± SD | 29,9 ± 4,4 | 24,8 ± 3,3 |  |  |  |  |  |  | 37,05 ± 0,23 | 36,94 ± 0,26 | 6,76 ± 0,22 |

Prior to uterine temperature measurement, the core body temperature of each woman was determined using a similar disposable temperature catheter (81-020409EU; DeRoyal) placed in the vagina. The mean core body temperature was 37.08 ± 0.26°C.

### Uterine temperature

We determined the temperature in the uterine cavity of 58 participants (51 unstimulated and 7 stimulated women; Table 1 and Figure 1a). The mean temperature in the human uterine cavity on the third day following a positive LH test or ovum pick-up was 36.94 ± 0.26°C (temperature range: 36.0°C to 37.6°C). One stimulated participant, participant 34, felt discomfort during temperature catheter insertion and therefore temperature measurement and further study procedures were aborted. The uterine temperature correlated with body temperature (r = 0.45, p < 0.001). The uterine temperature did not significantly differ between unstimulated women (36.94 ± 0.27°C) and stimulated women (36.99 ± 0.15°C) (ΔT = 0.05, p = 0.64).

**Figure 1.**
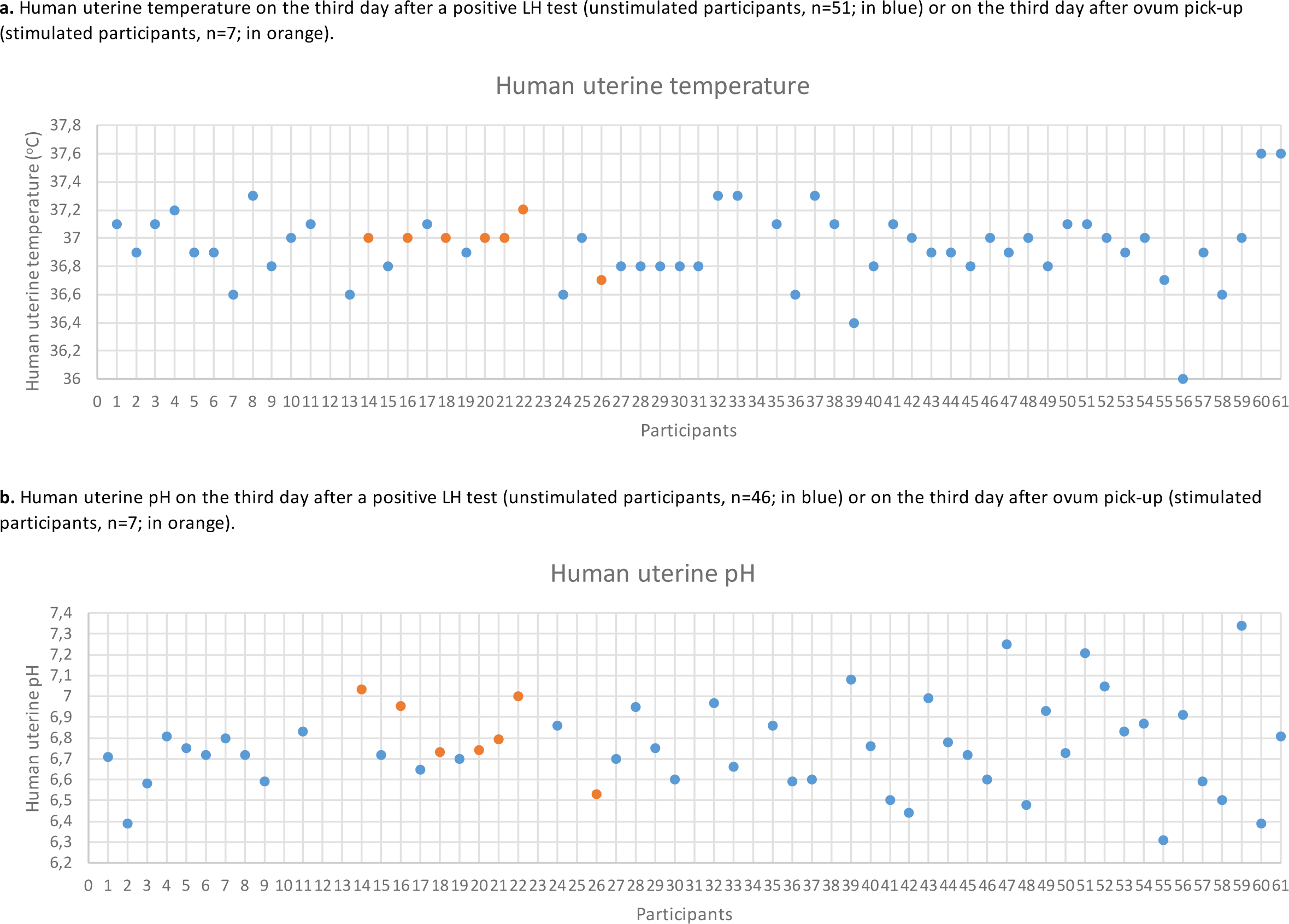
Human uterine temperature and pH.

### Uterine pH

We determined the pH in the uterine cavity of 53 participants (46 unstimulated and 7 stimulated women; Table 1 and Figure 1b). The mean pH in the human uterine cavity on the third day following a positive LH test or ovum pick-up was pH 6.76 ± 0.22 (pH range: 6.31 to 7.34). Five participants felt discomfort during temperature or pH measurement and therefore pH was not determined. The uterine pH did not significantly differ between unstimulated women (pH 6.75 ± 0.22) and stimulated women (pH 6.82 ± 0.18) (ΔpH = 0.07, p = 0.42).

### The composition of uterine fluid

We aspirated uterine fluid from 50 participants: 43 unstimulated and 7 stimulated women (Table 1 and Figure 1b). For practical reasons, the first available twenty (two) samples of aspirated uterine fluid were used to determine the mean concentrations of thirty-seven components of uterine fluid aspirated on the third day following a positive LH test (n = 15) or ovum pick-up (n = 7) (Table 2). This subset of components aligns with the thirty-seven components previously analysed in fifteen commercial human embryo culture media (Tarahomi et al. 2019). Important energy sources for preimplantation embryos were present in the following mean concentrations: 0.08 ± 0.02 mM pyruvate, 5.1 ± 1.78 mM glucose and 6.60 ± 1.12 mM lactate. The amino acids glutamic acid, glycine and alanine were present at much higher concentrations than the concentrations of other amino acids determined in the aspirated uterine fluid samples. Glutamic acid alone represented 23% of the total amino acid concentration in human uterine fluid. Citrulline, asparagine, tryptophan and methionine were present at the lowest concentrations: approximately 45-fold lower than glutamic acid. Furthermore, immunoglobulins were present in different concentrations, with IgG being the most predominant.

**Table 2.** Concentrations of thirty-seven components of human uterine fluid aspirated on the third day after a positive LH test (n=15) or on the third day after ovum pick-up (n=7). Concentrations are presented as the mean ± SEM.

| **Inorganic ions (mM)** |  | **Amino acids (μM)** |  |
| --- | --- | --- | --- |
| Calcium | 1.55 ± 0.18 | ***Essential amino acids*** | |
| Chloride | 82.34 ± 3.62 | Histidine | 97 ± 17 |
| Magnesium | 1.83 ± 0.17 | Isoleucine | 114 ± 14 |
| Phosphate | 3.30 ± 0.59 | Leucine | 193 ± 31 |
| Potassium | 17.22 ± 2.07 | Lysine | 298 ± 53 |
| Sodium | 143.81 ± 8.02 | Methionine | 38 ± 9 |
| | | Phenylalanine | 100 ± 15 |
| | | Threonine | 117 ± 20 |
| | | Tryptophan | 37 ± 4 |
| | | Valine | 190 ± 26 |
| **Energy sources (mM)** |  | ***Non-essential amino acids*** |  |
| Glucose | 5.1 ± 1.78 | Alanine | 513 ± 82 |
| Lactate | 6.60 ± 1.12 | Arginine | 136 ± 40 |
| Pyruvate | 0.08 ± 0.02 | Asparagine | 33 ± 7 |
| | | Aspartic acid | 230 ± 29 |
| | | Citrulline | 26 ± 3 |
| | | Glutamic acid | 1162 ± 183 |
| | | Glutamine | 227 ± 40 |
| | | Glycine | 955 ± 156 |
| | | Ornithine | 125 ± 10 |
| | | Proline | 221 ± 33 |
| | | Serine | 220 ± 46 |
| | | Tyrosine | 86 ± 15 |
| **Proteins (g/L)** |  |  |  |
| Albumin | 12.02 ± 1.23 | | |
| Total protein | 41.93 ± 4.24 | | |
| **Immunoglobulins (g/L)** |  |  |  |
| IgA | 0.42 ± 0.11 | | |
| IgG | 3.49 ± 0.44 | | |
| IgM | 0.43 ± 0.07 | | |
| **Other** |  |  |  |
| Iron (μM) | 115.11 ± 31.76 | | |
| Uric acid (mM) | 0.14 ± 0.03 | | |

## Discussion

*In vivo* evidence-based optimization of the *in vitro* embryo culture environment may improve human preimplantation embryo development and quality, and hold the potential to increase cumulative live birth rates and improve child outcomes after IVF/ICSI. However, the *in vivo* evidence on the natural environment of human preimplantation embryos is scarce. Better understanding of the *in vivo* characteristics of the preimplantation embryo environment in humans is essential to enable mimicking of this natural environment in an *in vitro* human embryo culture system. We aimed to gain fundamental knowledge on the human uterine conditions during the physiologically relevant period of the menstrual cycle. We measured the temperature (36.94 ± 0.26°C) and pH (6.76 ± 0.22) in the uterus of healthy women of reproductive age with normal ovulation and uterine function on the third day following a positive LH test or ovum pick-up, and determined the concentrations of thirty-seven components of uterine fluid aspirated at the same time in the menstrual cycle.

The strength of this study lies in the aim to analyse the characteristics of the human uterine cavity specifically at the physiologically relevant period of the menstrual cycle – when a preimplantation embryo normally resides and develops within the human reproductive tract. Although some previous studies have stated that *in vivo* measurement results throughout the menstrual cycle do not significantly differ from each other (Kermack, et al., 2015, Utsunomiya, et al., 2022), other studies have shown that uterine characteristics could change during the menstrual cycle (Feo, 1955, Yedwab, et al., 1976, Gardner, et al., 1996), possibly in response to shifting hormone levels, endometrial changes, or other factors. Since we want to use *in vivo* insights for optimization of *in vitro* culture environment for preimplantation embryos, it is important to ensure that the examined environment represents the natural preimplantation embryo environment as close as possible at the time of the measurements. We therefore timed the uterine measurements and uterine fluid aspiration in unstimulated women specifically on the third day following a positive LH test. However, the LH test yields a positive result at the onset of the LH surge, serving as an indicator of the impending ovulation rather than confirming its occurrence. Ovulation is anticipated 24-48 hours after a positive LH test result and therefore our analyses of uterine characteristics were conducted approximately 1.5 days post-ovulation. At this moment in the menstrual cycle, a zygote or preimplantation embryo could be hypothesised to still be in one of the oviducts. The time of arrival of an human embryo in the uterus is not established exactly, but generally assumed to be around the third day after fertilization, although it could be earlier or later. Regarding our timing, we may have identified an environment particularly suitable to cleavage stage embryos, which now could be considered a limitation. However, the measurements on the third day following ovum pick-up, approximately 3-3.5 days after ovulation in stimulated women – when an embryo is expected to reside in the uterus –, did not clearly differ from the results in unstimulated women. We also consider measurements in the uterus relevant for determining conditions for *in vitro* cleavage stage culture, because the timing of embryos entering the uterus has not exactly been determined yet, as we mentioned earlier, but mostly since measuring oviduct conditions comes with difficulties that challenge the results. Measuring oviductal pH is particularly difficult due to the narrowness of the oviducts, complicating access and accurate measurements. The use of CO_2_ for insufflation during surgeries alters the pH, a change that also occurs in ex vivo measurements following ovariectomy. Additionally, women scheduled for surgeries like ovariectomy often have reproductive health issues that may modify the oviductal environment.

Other strengths of this study include the extensive number of uterine measurements conducted at one specific point in the menstrual cycle. The reliability of the measurements is ensured by the use of catheters designed for internal temperature and pH measurements. Furthermore, we identified the concentrations of the largest number of components in each uterine fluid sample to date. Additionally, the reliable techniques used for determining the concentrations of the uterine fluid components add robustness to the results.

Limitations mainly regard the implications of the results. Firstly, a general and inevitable limitation of identifying the *in vivo* preimplantation embryo environment is the absence of a preimplantation embryo, which normally may have influenced its surroundings. Secondly, our study was confined to a single centre and a specific population of women. The potential influence of genetic or nutritional factors possibly limit the generalizability of the findings. Thirdly, uterine fluid likely consists of more components than the thirty-seven that we analysed. Fourthly, exact mimicking of all known characteristics of the *in vivo* environment *in vitro* may not provide an optimal human preimplantation embryo environment *in vitro*. Follow-up research on the effect of *in vivo* characteristics in an *in vitro* culture system for human preimplantation embryo culture will be necessary.

### Temperature

The mean uterine temperature we determined, 36.94 ± 0.26°C, was similar to the average temperature Yedwab et al. found in the uterine cavity of ten healthy women during their estimated ovulatory phase: 37.08 ± 0.18°C (Yedwab, et al., 1976). In absence of other *in vivo* studies, our findings add confidence to the suggestion that a temperature of 37.0°C is optimal for human embryo culture. The standard temperature of 37.0°C used in laboratory practice for human embryo culture in IVF matches these results.

### pH

The mean uterine pH we identified (pH 6.76 ± 0.22) slightly differed from the mean pH reported by two studies that previously investigated uterine pH (Feo, 1955, Sedlis, et al., 1967, Yedwab, et al., 1976). The early secretory pH values measured by Feo *et al*. (mean pH: 6.55, n=4; pH range 6.5-6.6) and early secretory and mid-secretory pH values measured by Sedlis *et al*. (mean pH: 6.5, n=17; pH range: 6.2-7.0) were a little lower, the mean pH reported by Yedwab *et al*. (pH 7.12, n=40) was higher. Despite several limitations, the results from our study and these other *in vivo* studies indicate that the pH of the human preimplantation embryo environment *in vivo* is lower than the pH currently considered optimal and widely used for *in vitro* human embryo culture (pH 7.2-7.4). These findings suggest that decreasing the pH of the *in vitro* human embryo culture system may improve human embryo development. Based on our study, we propose that *in vitro* embryo culture at a pH of 6.8 ± 0.1 may be more beneficial for human preimplantation embryos than the currently standard used pH of 7.3 ± 0.1. Future research should be conducted to confirm this.

### Uterine fluid composition

The amino acid concentrations we determined in the aspirated uterine fluid samples were fairly comparable to the concentrations of the 18 amino acids previously determined in 56 samples of human uterine fluid, although the latter were predominantly aspirated from subfertile women (Kermack, et al., 2015). Interestingly, amino acid concentrations that were recently identified in 28 samples of human midcycle and luteal oviduct fluid from infertile women were also similar (Utsunomiya, et al., 2022). Other components identified in these oviduct fluid samples, such as ions and energy sources (pyruvate, glucose and lactate), showed slightly diverse results with the concentrations of the components in our uterine fluid samples. For example, the average pyruvate concentration was higher (0.18 mM ± 1.48 mM) and the average lactate concentration was lower (4.66 ± 2.78 mM) than we determined in uterine fluid, but not significantly different. The average glucose concentration in midcycle oviduct fluid was lower (3.41 ± 1.61 mM) and in luteal oviduct fluid higher (4.16 ± 2.01 mM) than what we found in uterine fluid. Interestingly, the average magnesium concentration in oviduct fluid (0.53 ± 0.25 mM) appeared to be significantly lower than in uterine fluid. This difference seems relevant for embryo culture medium development, where two approaches (low or high magnesium concentration) have been used (Morbeck 2017).

One of the two small and older studies that had previously analysed uterine fluid composition, determined proliferative, midcycle and luteal phase uterine glucose, calcium, chloride, potassium, sodium, urea and fructose concentrations (Casslen and Nilsson, 1984). The midcycle and luteal phase mean glucose concentrations of respectively 5.7 mM and 5.2 mM (in total n = 6), align with the mean glucose concentration of 5.1 ± 1.78 mM in our samples. The mean concentrations of the analysed inorganic ions showed a similar pattern to our results, but were higher (potassium 25.8 mM and chloride 110 mM) or lower (sodium 110 mM) in absolute numbers. The other study, from 1996, investigated uterine concentrations of the metabolites pyruvate, lactate and glucose throughout the menstrual cycle (Gardner, et al., 1996). The mean glucose concentration (in total n = 15) was lower than 5 mM, namely 3.15 ± 0.31 mM. The mean concentrations of pyruvate (0.10 ± 0.05 mM) and lactate (5.87 ± 1.19 mM) were similar to our results.

In conclusion, our study contributes the most comprehensive analysis of human *in vivo* uterine temperature (n=58), pH (n = 53) and uterine fluid composition to date (37 components in 22 samples). We gained novel fundamental insights into the natural *in vivo* environment of human preimplantation embryos, valuable for assessment and optimization of the *in vitro* embryo culture system. The mean uterine temperature we found confirmed the use of 37.0°C for human embryo culture to be optimal for human preimplantation embryos. However, we observed a significantly lower pH (pH 6.76 ± 0.23) compared to the current widely adopted pH range of 7.2-7.4 for human embryo culture in IVF laboratories. Our results suggest that decreasing the pH of human embryo culture medium may be beneficial for the development and quality of human preimplantation embryos and potentially leads to a higher (cumulative) live birth rate. We therefore propose to test the effects of culturing human embryos in a pH of 6.8 ± 0.1. Furthermore, we provided a comprehensive overview of the concentrations of 37 components of uterine fluid, which can also be used for *in vivo* evidence-based embryo culture medium development.

## Data availability

The data underlying this article are available in the article.

## Authors’ roles

M.T., S.R. and S.M. designed the study. M.T., S.Z., S.F., A.M., Z.F., H.S and F.F. contributed substantially to data collection. M.T., J.v.S., F.S. and F.V. performed the uterine fluid analyses. M.T., M.Z. and M.v.W. analyzed and interpreted the data. M.Z., M.T., G.H. and S.M. wrote the final manuscript. All authors revised the manuscript and approved publication of the last version.

